# YAbS: The Antibody Society’s antibody therapeutics database

**DOI:** 10.1101/2025.02.07.637087

**Authors:** Puneet Rawat, Silvia Crescioli, R. Prabakaran, Divya Sharma, Victor Greiff, Janice M. Reichert

## Abstract

Therapeutic antibodies have gained prominence in recent years due to their precision in targeting specific diseases. As these molecules become increasingly essential in modern medicine, comprehensive data tracking and analysis are critical for advancing research and ensuring successful clinical outcomes. YAbS, The Antibody Society’s Antibody Therapeutics Database, serves as a vital resource for monitoring the development and clinical progress of therapeutic antibodies. The database catalogues detailed information on over 2,900 commercially sponsored investigational antibody candidates that have entered clinical study since 2000, as well as all approved antibody therapeutics. Data for the late-stage clinical pipeline and antibody therapeutics in regulatory review or approved (over 450 molecules) are openly accessible (https://db.antibodysociety.org). Antibody-related information includes molecular format, targeted antigen, current development status, indications studied, and the clinical development timeline of the antibodies, as well as the geographical region of company sponsors. Furthermore, the database supports in-depth industry trends analysis, facilitating the identification of innovative developments and the assessment of success rates within the field. This resource is continually updated and refined, providing invaluable insights to researchers, clinicians, and industry professionals engaged in antibody therapeutics development.

## Introduction

Natural human immunoglobulin (Ig), i.e., antibodies produced in the body, are glycoproteins composed of two identical heavy chains and two identical light chains that assemble to form a Y-shaped structure. Once assembled, these proteins have three key domains, two antigen-binding fragments (Fab) and one crystallizable fragment (Fc). This structure enables the antibody to carry out multiple functions, such as binding antigen via the Fabs and mediating effector functions via binding Fc receptors on cells such as natural killer cells, macrophages, neutrophils, and mast cells. [1] Over the past ∼ 40 years, the biopharmaceutical industry has incorporated these natural functions into recombinant monoclonal antibody (mAb) therapeutics.[2,3] Of the five classes of antibodies produced by human B cells (IgG, IgM, IgE, IgA, IgD), IgG (particularly the subclasses IgG1, IgG2, and IgG4) are the most common type of antibody therapeutic, although IgG3, IgM, and IgE mAbs have also been investigated in clinical studies.

Most currently marketed recombinant monoclonal antibodies have a canonical Y-shaped format. The investigational clinical pipeline, however, shows a clear trend toward the development of therapeutic antibodies that are more complex. [4–6] In addition to full-length canonical antibodies, the pipeline increasingly includes bi- or multispecific antibodies [7–10] and antibody-drug conjugates (ADCs), [11] as well as antibody fragments, fragment-Fc and appended Ig, which can be naked, conjugated to other molecules or fused to non-Ig proteins, and can have monospecific, bispecific or multispecific properties.

In an ongoing effort to support individuals and organizations involved in antibody therapeutics research and development, The Antibody Society, Inc., an international non-profit trade association, provides business intelligence on global antibody therapeutic development conducted by the commercial sector. As part of their research, the Society determines development trends, success rates (clinical phase transition rates and overall marketing approval rates), [5,6,12] and clinical development phase lengths for antibody therapeutics, which are key metrics that inform decisions about resource allocation by the biopharmaceutical industry. Given the increasingly complex nature of therapeutic antibodies and the substantial growth of commercial antibody therapeutics development in the past decade, [4,5] a comprehensive resource for efficient tracking of antibody candidates is essential to The Antibody Society’s business intelligence research. Several public (e.g., IMGT/mAb-KG, [3] DrugBank [13] TheraSAbDab [14] and commercial (e.g., Beacon (beacon-intelligence.com)) databases are available, but none adequately meet our specific objectives due to the absence of key clinical, regulatory, and business development milestone data or insufficient comprehensiveness. To address this issue, the Society has developed and maintains their own dataset, which currently includes data for over 2,900 commercially sponsored antibody therapeutics at various stages of development. The dataset enables a broad assessment of the global biopharmaceutical industry’s pipeline. Analyses can be performed on antibody therapeutics as a class, as well as on particular aspects, such as format (e.g., full-length antibody, fragment), targeted antigen, therapeutic area, development status (e.g., preclinical, Phase 1, regulatory review), or geographic region of the sponsors.

The Society extensively analyzes the data and delivers up-to-date insights via presentations and reports, such as the “Antibodies to Watch” article series published in *mAbs* (tandfonline.com/journals/kmab20/collections/antibodies-to-watch), which informs and educates stakeholders interested in antibody therapeutics development. Extracts of the dataset, covering over 220 antibody therapeutics that are approved or currently in regulatory review (https://www.antibodysociety.org/antibody-therapeutics-product-data/) and over 170 molecules in late-stage clinical development (https://www.antibodysociety.org/antibodies-in-late-stage-clinical-studies/), are available in searchable tables on The Antibody Society’s website.

Here, we introduce YAbS, a database developed to provide a web interface for searching, filtering, analyzing, and exporting antibody therapeutics data. YAbS offers extensive filtering and search options based on standardized nomenclature, functionality and architecture for variables such as molecular category and format, target antigen, development status, therapeutic area, company sponsor, and country of origin. Furthermore, we provide three use cases, highlighting the utility of the database as a categorical stratification or trends over time assessment. The YAbS database, accessible at https://db.antibodysociety.org, currently offers open access to data for over 450 antibody therapeutics that are approved, in regulatory review, or in late-stage clinical development. Overall, YAbS is a standardized antibody therapeutics resource for tracking past and upcoming clinical candidates along with their developmental histories. It offers crucial insights to support decision-making by researchers, clinicians and industry professionals engaged in antibody therapeutic development.

### Decoding the YAbS database and antibody therapeutic data

The YAbS database was developed in Python 3.9 using the Django framework, with the dataset stored in an SQL database, as previously described [15,16]. It serves as the searchable interface for a dataset of antibody therapeutics that was created in 2012 and subsequently updated regularly. We have established strict criteria for molecules that are included in the database. Specifically, the molecule must be a novel, therapeutic, recombinant protein with at least one antigen-binding site derived from an antibody gene, developed or in-licensed by a company, with initial clinical entry on or after January 1, 2000. Exceptions are made for approved antibody therapeutics that began clinical studies in the 1980s or 1990s and molecules in late preclinical development. By definition, the following categories of molecules are excluded: 1) Biosimilar antibodies; 2) Antibodies used solely for non-therapeutic purposes, such as disease diagnosis; 3) Polyclonal antibodies derived from a natural source; 4) Related molecules that are not protein or do not contain an antibody-derived binding site, such as antibody-encoding DNA, alternative targeted proteins (e.g., designed ankyrin repeat proteins, Anticalin® proteins), and Fc fusion proteins (i.e., proteins containing only the Fc portion of an antibody); 5) Antibody therapeutics in clinical studies sponsored solely by non-commercial organizations and not included in a company pipeline; 6) Antibody therapeutics that entered clinical study prior to January 1, 2000, unless they were granted a marketing approval.

The Antibody Society continuously collects top-level data for antibody therapeutics from public sources, including company websites, press releases and presentations, clinical trials registries, regulatory agencies, the World Health Organization’s lists of recommended international non-proprietary names (INN), open-access databases of therapeutic antibodies such as TheraSAbDab and IMGT/mAb-DB, as well as literature reports and reviews. The YAbS database is updated bimonthly from data collected daily, to track the development of antibody therapeutics, from preclinical development to regulatory approval.

To transition from the dataset to a structured database, we first standardized the nomenclature to annotate antibody therapeutics. The format category encompasses full-length antibodies, antibody fragments with or without an Fc, and appended Igs. Fragments may contain one or more Ig domains capable of antigen binding (e.g., VH, VL, VHH, Nanobody), scFv, dsFv, Diabody, DART, Fab, F(ab)2). Fragment-Fc are composed of one or multiple antibody fragments fused to an Fc, while appended Ig are full-length Ig fused to one or multiple antibody fragments. Each of these formats can be naked, conjugated to other molecules such as small-molecule drugs or fused to non-Ig proteins (**Supplemental Figure 1**). We annotated the general format of the molecules according to the schematic in **Supplemental Figure 1** and the format detail according to the schematic in **Supplemental Figure 2**. We further categorized therapeutic antibodies using the following features: specificity, conjugation status, composition (i.e., one molecule or a mixture of two or more), mechanism of action, and antigen binding properties. A therapeutic antibody can be monospecific, bispecific, or multispecific. If bispecific or multispecific, we noted whether it binds to different antigens or different epitopes on the same antigen (biparatopic or multiparatopic). Except for ADCs, and Radioimmunoconjugates, we classify antibodies that are conjugated to non-protein molecules or fused to non-Ig proteins or protein domains as Immunoconjugate. Among immunoconjugates, we also included molecules which fall under the unconventional ADC category, such as Antibody Degrader Conjugate, Immune-Stimulating Antibody Conjugate, Antibody Oligomer Conjugate, Antibiotic Conjugate, and Steroid Conjugate. Furthermore, antibody therapeutics may be formulated as a single molecule or a mixture of molecules, have canonical mechanisms of action (blocking, agonist, antigen clearance, cell mediated effector function, payload delivery) or function as vaccines, or be engineered with canonical or conditionally active antigen binding properties. We used one or more of these categories to annotate the general molecular category of the molecules, as described in the schematic in **Supplemental Figure 3**.

As of January 2025, the database contains information for over 2900 antibody therapeutics, which can be searched in a variety of ways (**Figure 1**) to yield the desired output. The search options and result table are dynamically generated, allowing for future expansion based on new entries. Users can perform a quick search based on target, therapeutic area, or the company developing the molecule (**Figure 1A**), or they can use the advanced search page (**Figure 1B**). On the advanced search page, users can search based on the name of the molecule (INN or drug code), or apply filters related to the molecular characteristics and clinical development. Molecular characteristic variables include the general molecular category (e.g., ADC, bi- or multi-specific, immunoconjugate), target antigen, format (general format category e.g., full length, fragment, appended Ig), heavy and light chain isotype and sequence source. Clinical development variables include general status (e.g., preclinical, clinical, approved), or more specific status, such as the most advanced stage of development (e.g., Phase 1, Phase 2, Phase 3), and the primary therapeutic area of development. A dropdown menu allows users to select milestone clinical development events, while a slider enables date-based filtering to retrieve results for specific timeframes. Users can also search by the location of the company developing the molecule. A Free Text Search field allows the user to search for specific text in any column. To the best of our knowledge, the database is unique in allowing filtering by date (e.g., initiation of clinical study, Phase 2 or 3 start, first approval), which allows analyses of development trends over time. Users can filter based on one parameter or a combination of parameters. Schematics of the general format, detailed format, and general molecular category classification systems can be found in **Supplemental Figures S1, S2, and S3**, respectively. A complete list and descriptions of the database variables are provided in **Supplemental Table S1**.

**Figure 1.**
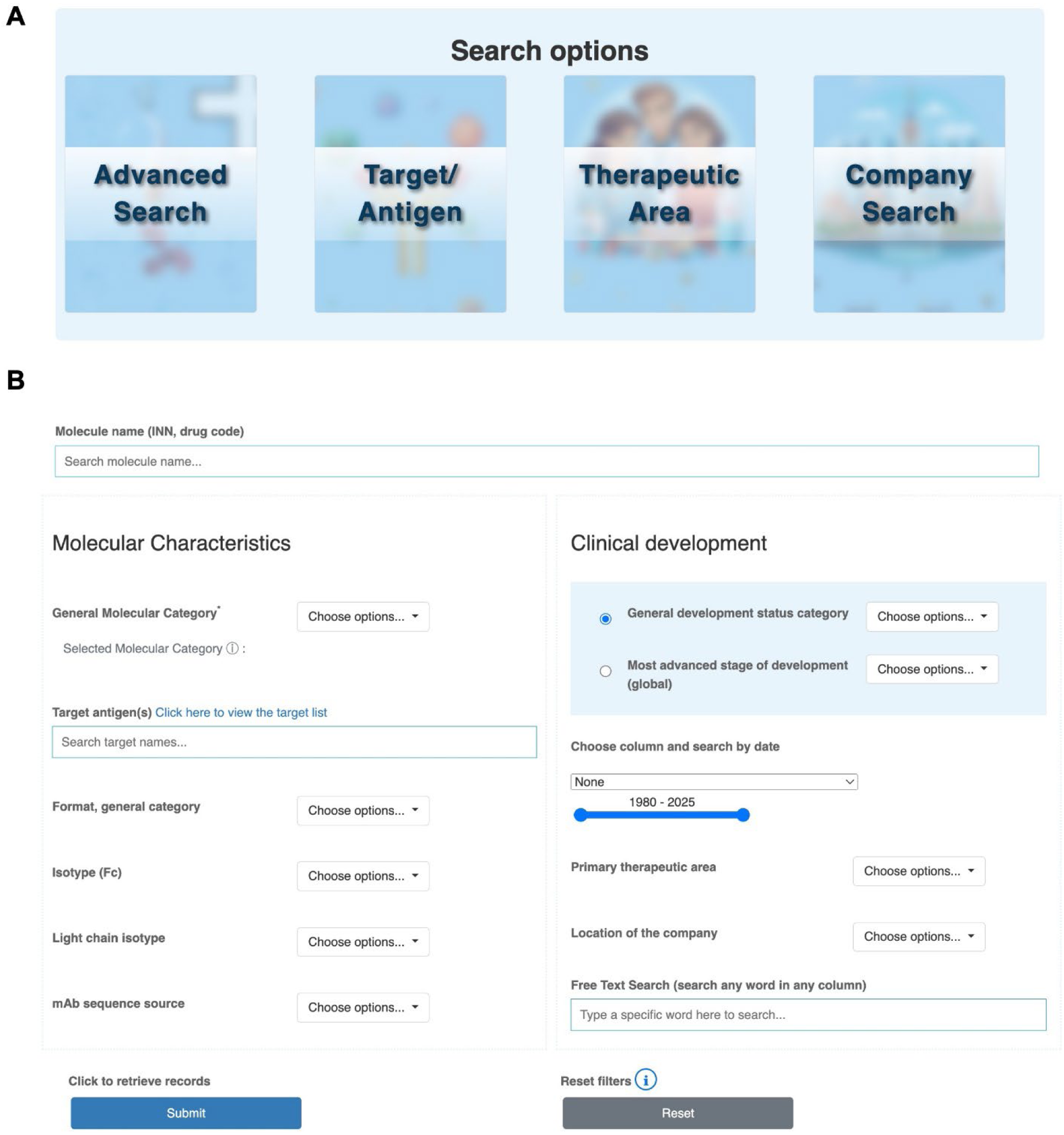
YAbs database search and filtering options. (A) The database homepage includes links to four different ways to search antibody molecules: three quick search pages based on target, therapeutic area, or name of the company developing the molecule, and an advanced search page (B) with multiple customizable filters.

The advanced search interface enables both broad searches (such as all antibodies currently in clinical studies), and highly specific queries by applying multiple filtering parameters (such as bispecifics targeting BCMA and CD3 that are in Phase 3 clinical studies for cancer). The search by time feature allows filtering based on milestone event dates, such as the start of the clinical study period, initiation of the first Phase 2 and Phase 3 study, submission dates for biologics license applications (BLAs), and first approval dates.

It should be noted that not all companies are forthcoming about the nature of the molecules they are developing, particularly those in preclinical and early-stage clinical development. Thus, at any given time, specific data about molecular characteristics, such as target, Fc and light chain isotype, and identity of any fused or conjugated component, may not be available for all antibodies in the database. Moreover, the indications or therapeutic area may not be known for molecules in early-stage clinical studies if the company does not divulge the information and the Phase 1 studies are conducted in healthy volunteers. As noted above, records for the molecules in the database are continuously updated as new information enters the public domain.

After performing a search using the advanced search panel, users are directed to a result page that displays the output in the form of pie charts and a table (**Figure 2**). The pie charts are created using the Chart.js library (https://www.chartjs.org/). Four pie charts represent the distribution of the general status, the general molecular category, the primary therapeutic area and the targets of the filtered molecules. Search boxes located at the top of the results page allow users to refine their search further by applying additional filtering based on development status, molecular category, therapeutic area, and target. The table provides detailed data for the filtered molecules, with customizable columns. Users can select which columns to display and visualize the additional columns by scrolling right. The table can be sorted based on a certain column, by clicking on the column header. The filtered records can be exported, and the exported table will include all columns for the filtered records.

**Figure 2.**
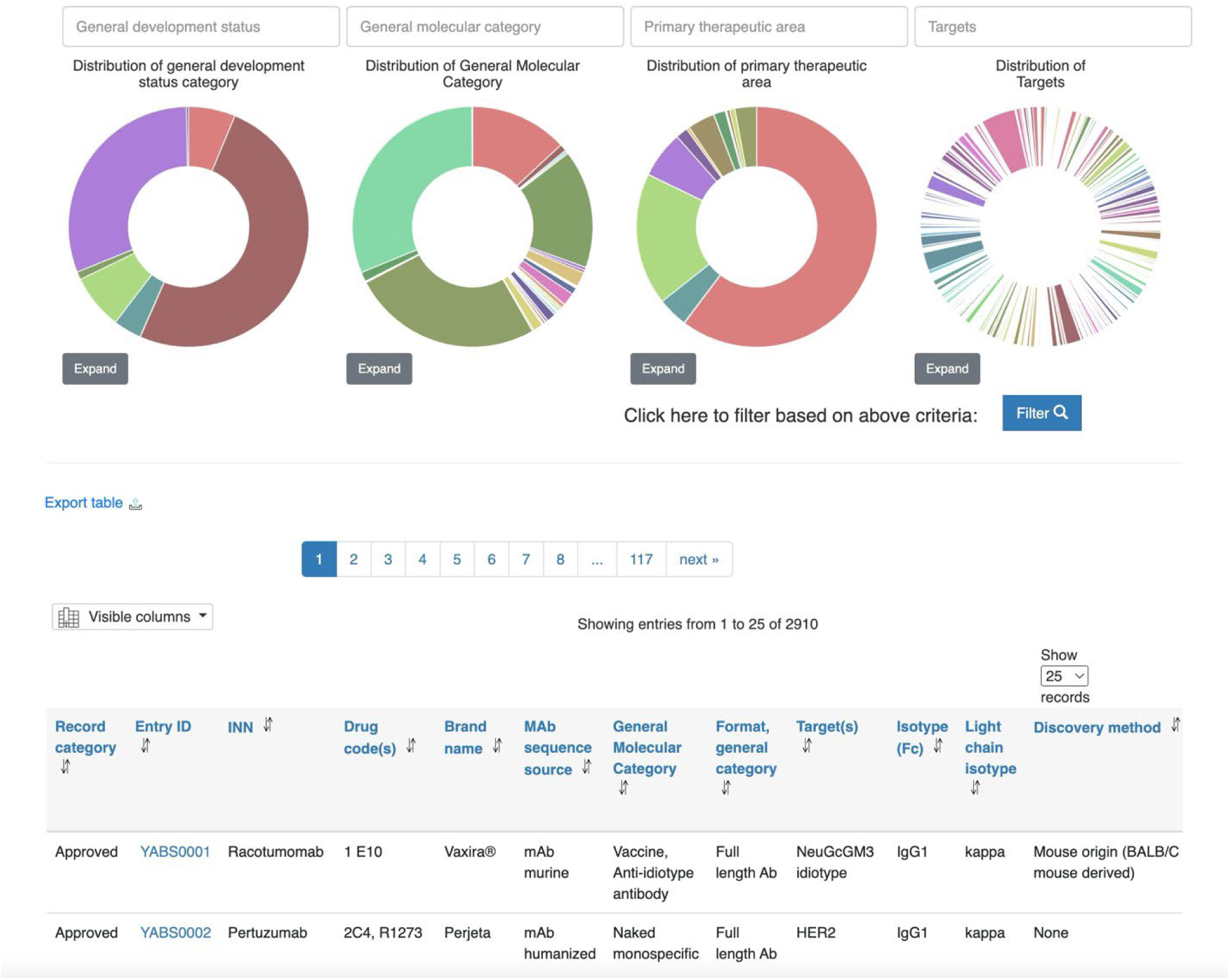
YAbs database results page: The results page is divided into two sections. The top section contains pie charts and corresponding filtering boxes, providing a quick graphical overview of the results and allowing further filtering based on general development status, general molecular category, primary therapeutic area, and target. The bottom section features a table displaying all filtered molecules, with options to select which columns to display, adjust the number of records per page, and the option of sorting the table by clicking on column headers. Additional selected columns on the table can be visualized by scrolling right. Clicking on the Entry ID provides detailed information about the antibody candidate. An additional button allows users to export the filtered records. The exported table will include all columns for the filtered records.

The Entry ID column displays the unique YAbS ID for each molecule. Users can view the dedicated page for each molecule by clicking on the molecule ID. Examples of molecule-dedicated pages can be found in **Supplemental Figure S4 and S5**. On the dedicated page, each molecule is described according to dataset parameters grouped as: 1) Antibody information, such as format and isotype; 2) Therapeutic information, such as target, indications studied, and primary therapeutic area; 3) Development information, such as the most advanced stage of development, key dates of clinical development, approvals, and development timelines; 4) Company information, which may include information on specific clinical trials, regulatory agency designations, and company acquisitions and collaborations, and the location of the originating company; 5) Description/comment, which may include summarized details and relevant publications; and 6) Additional information, which may include anticipated events, such as the start of a clinical study or submission of a marketing application, as well as reasons for termination of development.

### Using the database to evaluate key antibody therapeutics development trends

The YAbS database has multiple use cases. First, the database is the repository for top-level data on the current commercial clinical pipeline of antibody therapeutics, enabling real-time knowledge of company portfolios and upcoming events. Second, analyses of key variables (such as antibody format, target, and indication) allow the determination of trends in innovative antibody therapeutics development over time, findings that are regularly included in The Antibody Society’s presentations and published reports [4–6]. Third, the database, to the best of our knowledge, includes the most up-to-date status of all publicly disclosed, commercially sponsored antibody therapeutics that were first administered to humans after January 1, 2000, which enables the calculation of accurate success rates for these molecules. Several analyses of success rates derived from data in the YAbS database have been previously published [5,12]. Below, we discuss three examples of YAbS-derived data analyses.

### Use case 1: Assessment of clinical-stage molecules

**Figure 3** illustrates the results of four broad searches of the YAbS data for antibody therapeutics evaluated in clinical studies (i.e., preclinical molecules were excluded). The data was initially stratified by the top-level development status (**Figure 3A**). The molecules in active clinical development were then further stratified by current clinical phase, therapeutic area, and company region (**Figures 3B-D**). A detailed analysis pipeline describing the Advanced Search panel filters used for each graph is shown in **Supplemental Figure S6**. Our results show that the majority (55%) of these antibodies are in active clinical development (**Figure 3A**). Among them, most are in early-stage development, with nearly three-quarters in Phase 1 or 1/2 clinical studies, (**Figure 3B**) and the majority (66%) are treatments for cancer (**Figure 3C**). Notably, most of the molecules currently in clinical studies originated at companies based in China or the US (**Figure 3D**).

**Figure 3.**
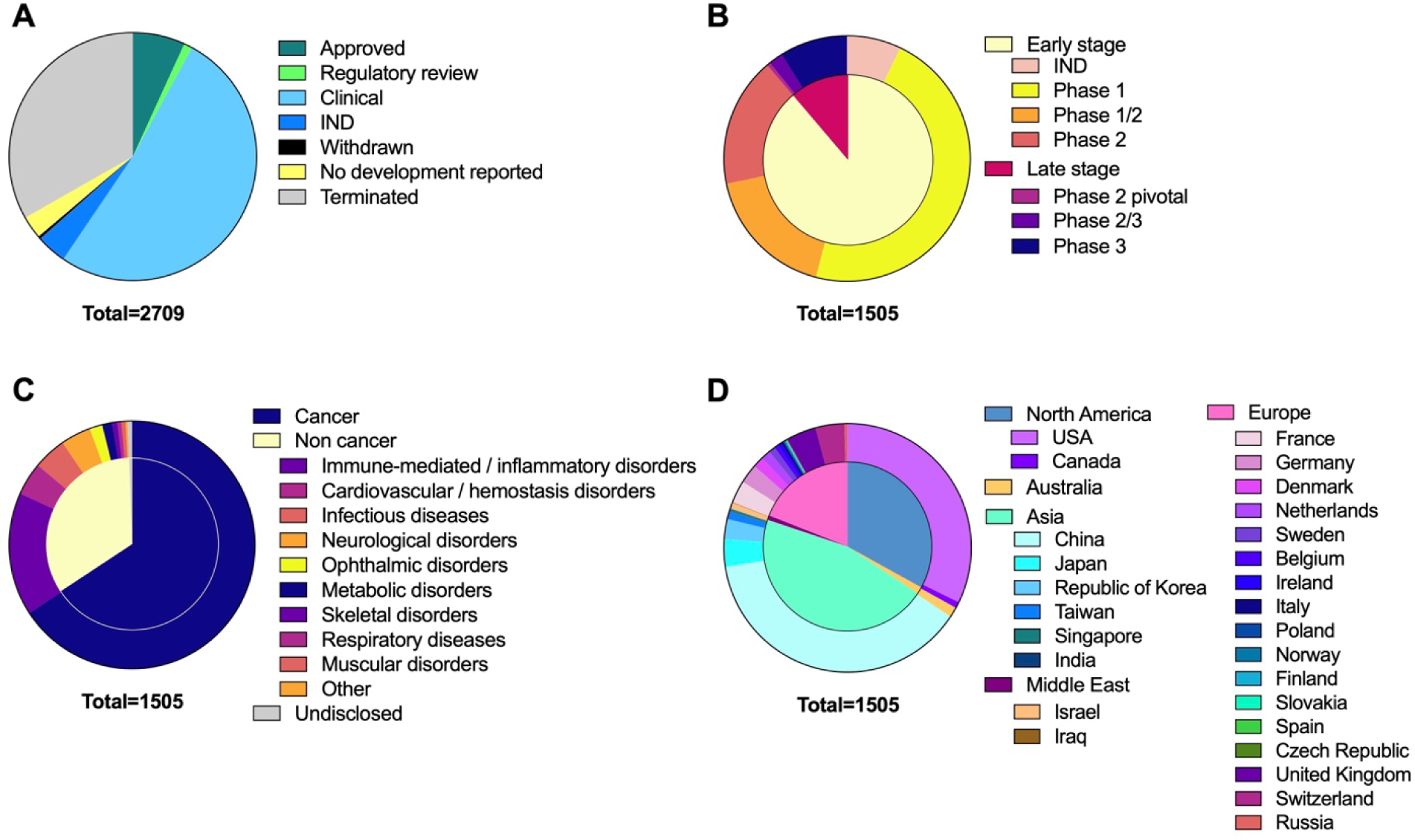
The current development status of antibody therapeutics that entered clinical study after January 1, 2000, stratified by clinical phase, therapeutic area, and company region. (A) Pie chart representing the development status of antibody therapeutics that entered clinical studies (n=2709). Data for the molecules currently in clinical studies (n=1505) was stratified by (B) most advanced stage of clinical development; (C) therapeutic area; and (D) region and country of the company developing the molecule. Data from the YAbS database as of December 12, 2024. A detailed analysis pipeline describing the Advanced Search panel filters used for each graph is shown in **Supplemental Figure S6**.

### Use case 2: Assessment of trends in antibody therapeutics development

Several examples of analyses designed to reveal trends in antibody therapeutics development over time are shown in **Figure 4**. A detailed analysis pipeline describing the Advanced Search panel filters used for each graph is shown in **Supplemental Figure S7**. Each panel shows an aspect of trends in annual first-in-human (FIH) studies of antibody therapeutics sponsored by commercial firms during 2010 to 2023. **Figure 4A** highlights the substantial increase in the number of annual FIH studies during this period and shows the current status of the molecules. While the overall trend clearly indicates greater participation in commercial antibody therapeutics development, downturns compared to the previous year can be observed in 2012, 2016, 2019, and 2023. Various factors, including budget re-assessments, changes in company priorities, and altered projections about the medical landscape, can influence company decisions to initiate FIH studies. The data also show subsequent increases in the annual number of FIH studies after downturns (i.e., in 2013, 2017, and 2020). We await further data to determine if the number of FIH studies started in 2024 will exceed those of 2023. As would be expected, molecules that started in clinical studies recently are more likely to still be in clinical studies compared to those that entered clinical studies earlier, which are more likely to be terminated or approved. **Figure 4B** presents the same annual FIH study data but categorizes molecules by general molecular type instead of current status. Substantial increases in the number of bi- and multi-specific antibodies and ADCs is evident in **Figure 4B**. Trends in the development of antibodies targeting the well-validated human epidermal growth factor-2 (HER2) antigen over the 2010-2023 period are shown in **Figure 4C**. For reference, we note that the naked, monospecific, anti-HER2 antibody trastuzumab (Herceptin®) was developed in the 1990s and granted a first marketing approval in 1998. Our analysis reveals that antibodies targeting HER2 were actively developed during 2010-2023, but the categories of the molecules have diversified. HER2 is now the target of ADCs, multispecifics, ADC multispecifics, immunoconjugates, and antibodies for radioimmunotherapy, in addition to naked monospecific antibodies.

**Figure 4.**
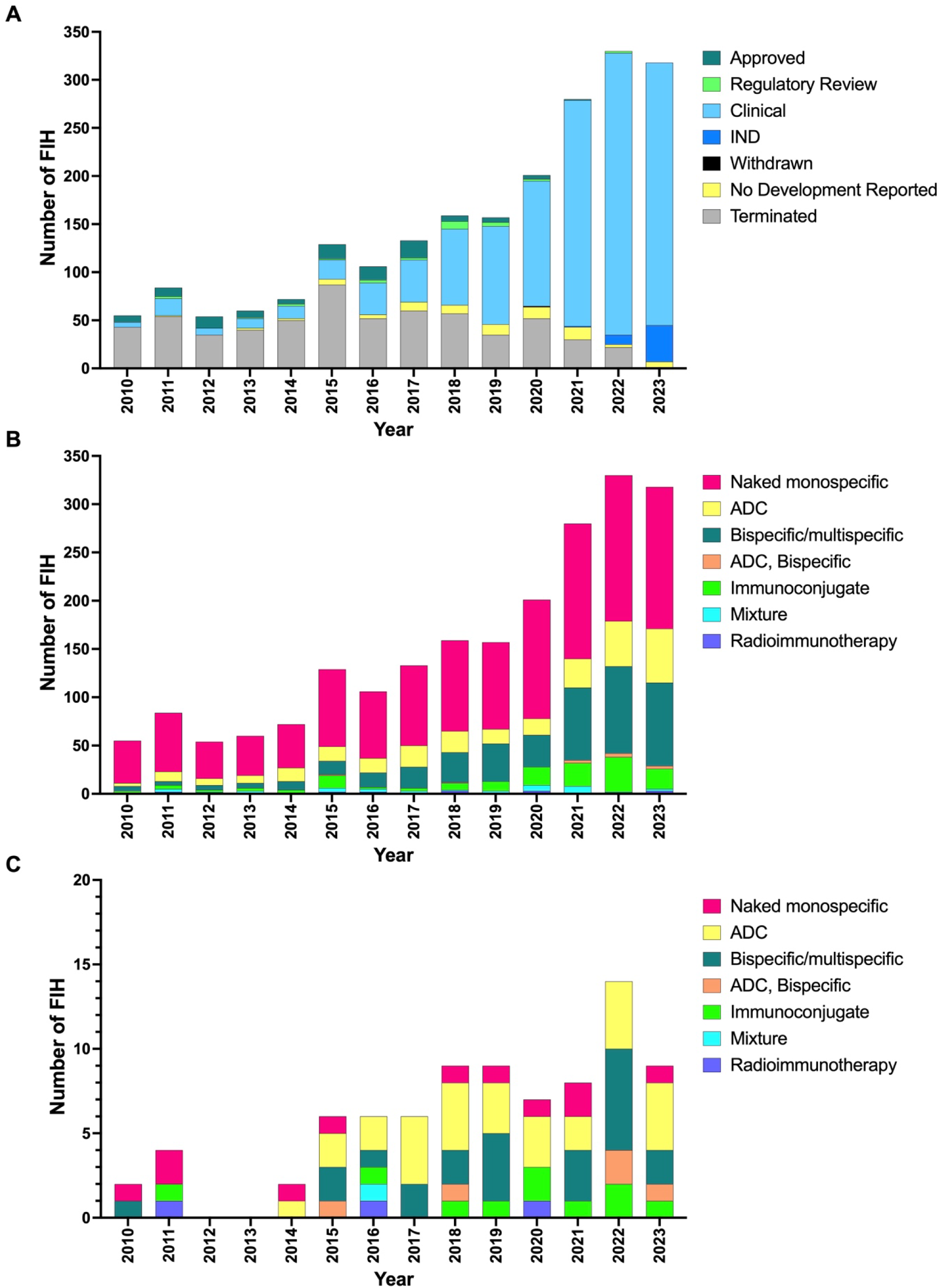
Annual number of first-in-human (FIH) studies of antibody therapeutics initiated between 2010 and 2023. (A) Annual number of FIH studies stratified by current status (n=2138), (B) Annual number of FIH studies stratified by general molecular category (n=2138), and (C) Annual number of FIH studies of anti-HER2 antibodies stratified by general molecular category (n=82). Data sourced from the YAbS database as of December 12, 2024. A detailed analysis pipeline, describing the advanced search panel filters used for each graph is shown in **Supplemental Figure S7**. Abbreviations: ADC, antibody-drug conjugate; FIH, first-in-human study.

### Use case 3: Assessment of milestone events for different therapeutic molecules

The results in **Figure 5** demonstrate another use for the milestone event dates. The inclusion of key dates for the start of first early- and late-stage studies, as well as BLA submission and US Food and Drug Administration (FDA) approval dates, enables the calculation of average phase lengths. A detailed analysis pipeline describing the Advanced Search panel filters used for each graph is shown in **Supplemental Figure S7**. The majority of antibody therapeutics entering clinical studies after January 1, 2000 and approved by the FDA by December 31, 2023 were treatments for cancer (**Figure 5A**). When stratified by general therapeutic area (cancer or non-cancer), we observed that, on average, the clinical development and approval period for antibodies developed for non-cancer indications was approximately 1 year longer overall due to longer late-stage clinical study and regulatory review periods (**Figure 5B**). In general, drugs for cancer are more likely to be granted FDA designations, such as Breakthrough Therapy, Fast Track and Priority review, that are designed to speed the clinical and regulatory review periods, which may be one reason for the differences. **Figure 5C** illustrates that, when further stratified by detailed therapeutic area, nearly half the antibodies approved for non-cancer indications are treatments for immune-mediated or inflammatory diseases. Phase lengths for the top four non-cancer indications show substantial variation, with cardiovascular/hemostasis agents having the shortest average total clinical and regulatory period (slightly less than 7 years) and immune-mediated or inflammatory disease agents having the longest (nearly 10 years) (**Figure 5D**). We stratified the data for FDA-approved antibodies for cancer indications by general molecular category and observed that just over half are naked, monospecific antibodies (**Figure 5E**). According to our phase length analyses, these types of antibodies have a longer average total clinical and regulatory period compared to ADCs and bispecific antibodies (**Figure 5F**). It should be noted that a relatively small number of ADCs and bispecifics have been approved, and thus these average phase lengths may change in the future.

**Figure 5.**
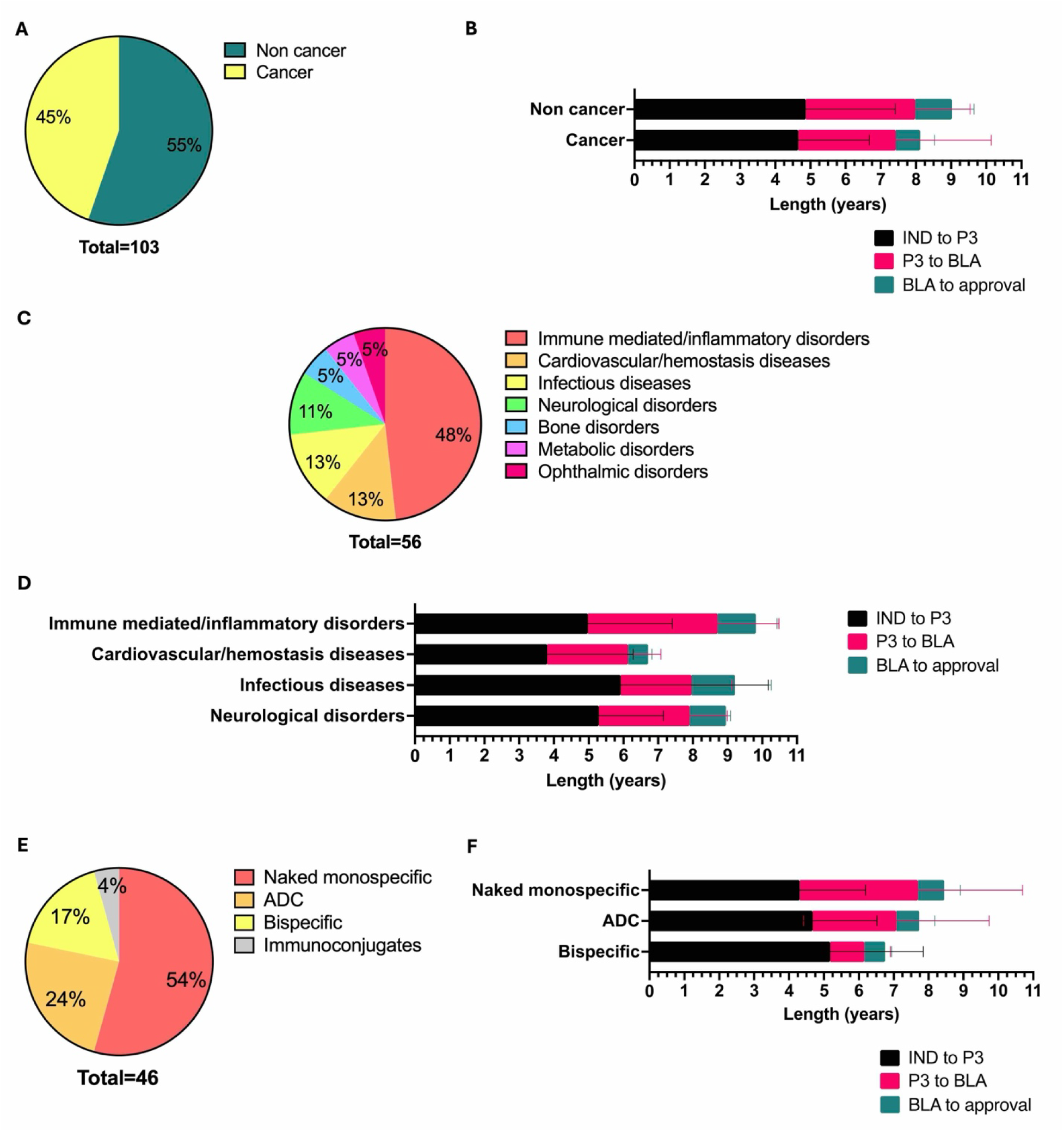
Development phase lengths of antibody therapeutics entering clinical studies after January 1, 2000 and approved by the US Food and Drug Administration by December 31, 2023. (A) Pie chart representing the percentage of approved antibodies for non-cancer and cancer indications (n=103). (B) Stacked bar charts representing mean phase lengths for antibodies for non-cancer and cancer indications. (C) Pie chart representing approved antibodies for non-cancer indications stratified by therapeutic area (n=56). (D) Stacked bar charts representing mean phase lengths for approved antibodies in the top four therapeutic areas for non-cancer indications (n=47). (E) Pie chart representing approved antibodies for cancer indications stratified by general molecular category (n=46). (F) Stacked bar charts representing mean phase lengths for approved antibodies in the top three general molecular categories for cancer indications (n=44). (B, D, F) Error bars represent standard deviation. A detailed analysis pipeline describing the Advanced search panel filters used for each graph is shown in **Supplemental Figure S7**. Data sourced from the YAbS database. Abbreviations: ADC, antibody-drug conjugate; BLA, biologics license application; IND, investigational new drug application; P3, first Phase 3 clinical study.

## Discussion

The YAbS database is a comprehensive and dynamic resource for researchers, clinicians, and industry professionals. It is designed to address the growing need for detailed and accessible information on therapeutic antibodies, offering insights into the molecular characteristics, developmental timelines, and clinical progress of over 2,900 antibody candidates. The creation of this resource reflects the increasing importance of antibody therapeutics in modern medicine and the challenges associated with their development. One of the key strengths of the YAbS database is its ability to track antibody therapeutics across various stages of development, from preclinical studies to marketing approvals, over time. This capability enables insights into the landscape of antibody therapeutics, such as identification of emerging trends, innovative developments, and potential gaps in the market. However, it is important to note that some information, particularly for antibodies in early-stage development, may not be fully disclosed by companies, leading to gaps in the data available at any given point in time. Despite this limitation, the database provides a robust foundation for understanding the current state of antibody therapeutics development. The inclusion of detailed molecular data, such as general molecular category, targets, formats, Fc and light chain isotypes, and conjugated components, allows in-depth analysis and comparison of different antibody candidates.

Furthermore, the database’s user-friendly interface and dynamic search options enable users to perform both broad and specific searches, making it a versatile tool for various research and development needs. For instance, users can perform a quick search for antibodies based on their target, therapeutic area, or location of the developing companies. For a more advanced search, users can filter the dataset based on the name of the molecule (INN or drug code), or use filters related to the molecular characteristics and clinical development of the antibody. The ability to filter results by time periods and milestone events, such as the start of clinical trials or submission of BLAs, provides valuable insights into the pace of development and regulatory progress of antibody therapeutics. The database allows users to both extract the filtered data and to find details on specific molecules. The dedicated page for each antibody candidate includes key information about clinical development (e.g., publicly disclosed upcoming events, dates of clinical transitions, clinical trial numbers) and the companies involved in development (e.g., company acquisitions and collaborations) that is not usually featured in public databases.

Insights derived from the YAbS database can have substantial implications for the future of antibody therapeutics. For example, the trends observed in FIH studies, as well as the development of specific molecular categories such as bispecifics and ADCs, highlight the evolving strategies in antibody design and application. The detailed analysis of phase lengths for antibodies developed for cancer versus non-cancer indications provides valuable information on the challenges and opportunities in different therapeutic areas. The ability of the YAbS database to support ongoing and future research is enhanced by its potential for expansion. As new antibody candidates are developed and enter clinical trials, the database will be updated to reflect these changes, ensuring that it remains a relevant and up-to-date resource.

In summary, the YAbS database is a powerful tool that enhances understanding of the antibody therapeutics landscape. By providing comprehensive, detailed, and dynamic data, it supports informed decision-making and fosters the advancement of therapeutic antibody research and development. The continued maintenance and expansion of this database will be crucial in keeping pace with the rapid developments in the field, [17,18] ensuring that it remains an invaluable resource for all stakeholders involved in antibody therapeutics development.

## Disclaimer

Data included in YAbS are collected from the public domain. The Antibody Society, Inc. makes no representation or warranties with respect to the accuracy, completeness, or timeliness of the information and specifically disclaims any implied warranties of fitness for a particular purpose. The Antibody Society, Inc. assumes no liability, contingent or otherwise, for the accuracy, completeness, or timeliness of the Information, or for any decision made or action taken in reliance upon the information. The Antibody Society, Inc. reserves the right to control access to the YAbS database. Data for the late-stage clinical pipeline, antibody therapeutics in regulatory review, and those granted marketing approvals in any country are freely available. Access to data for antibody therapeutics in early-stage development and those terminated while in development or after approval is currently limited.

## Abbreviations

Ab: antibody
ADC: antibody-drug conjugate
BLA: biologics license application
Fab: antigen-binding fragment
Fc: crystallizable fragment
FDA: US Food and Drug Administration
FIH: first-in-human
HER2: human epidermal growth factor-2
Ig: immunoglobulin
IND: investigational new drug
INN: international non-proprietary name
mAb: monoclonal antibody

## Supplemental Materials

**Supp. Figure S1.**
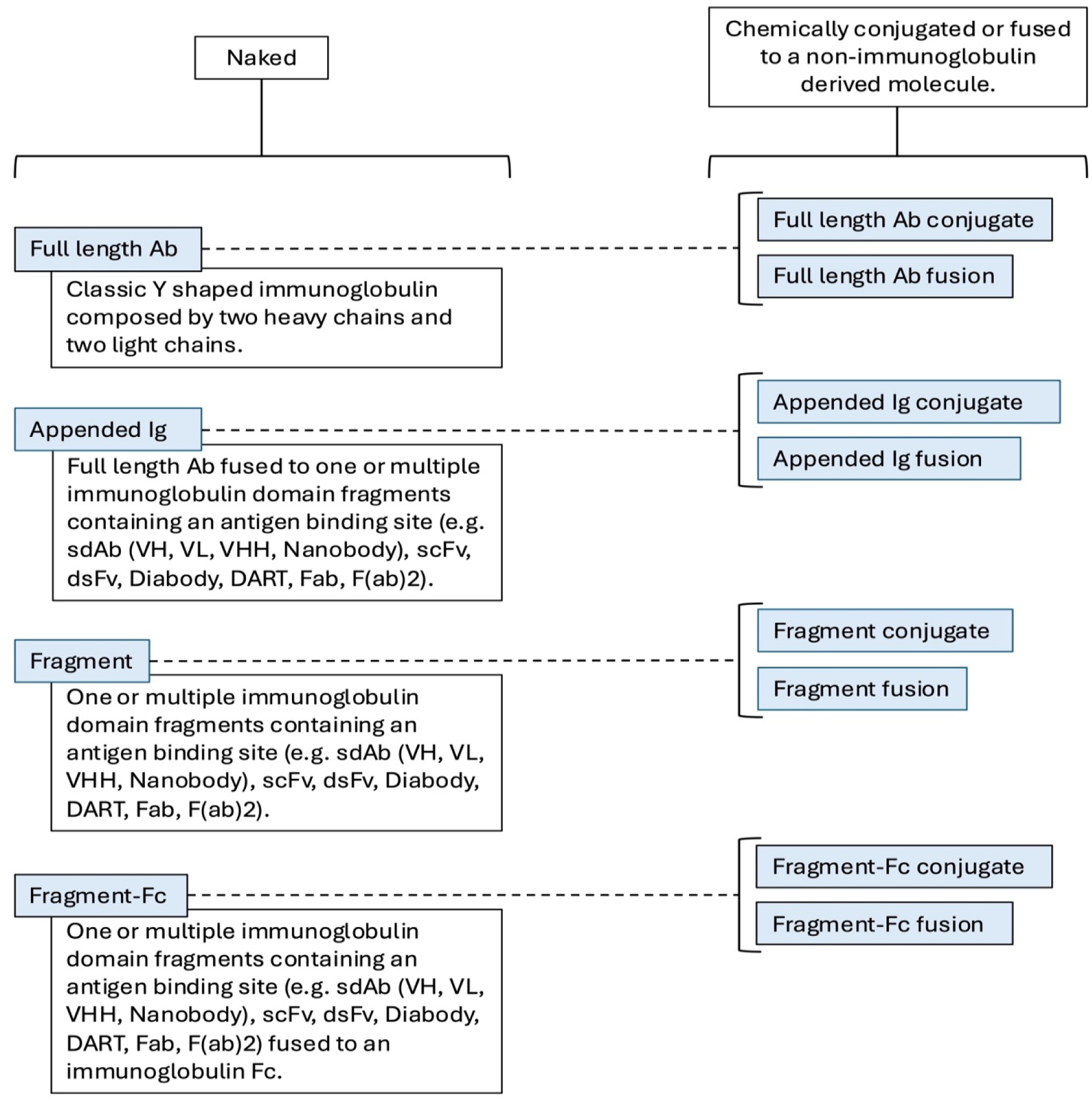
Schematic of general format classification system. The schematic represents the rationale behind our standardization method to describe the general format of therapeutic antibodies. We first divided them into two groups: naked antibodies and antibodies that are chemically conjugated or fused to a non-immunoglobulin derived molecule. We further divided naked antibodies into four categories as described in the schematic: Full length Ab, Appended IgG, Fragment, Fragment Fc. Each of these categories can also be conjugated or fused to one or more non-immunoglobulin derived molecules, as shown on the schematic. Blue boxes represent the variables represented in the database. Only one variable can be assigned to a molecule.

**Supp. Figure S2.**
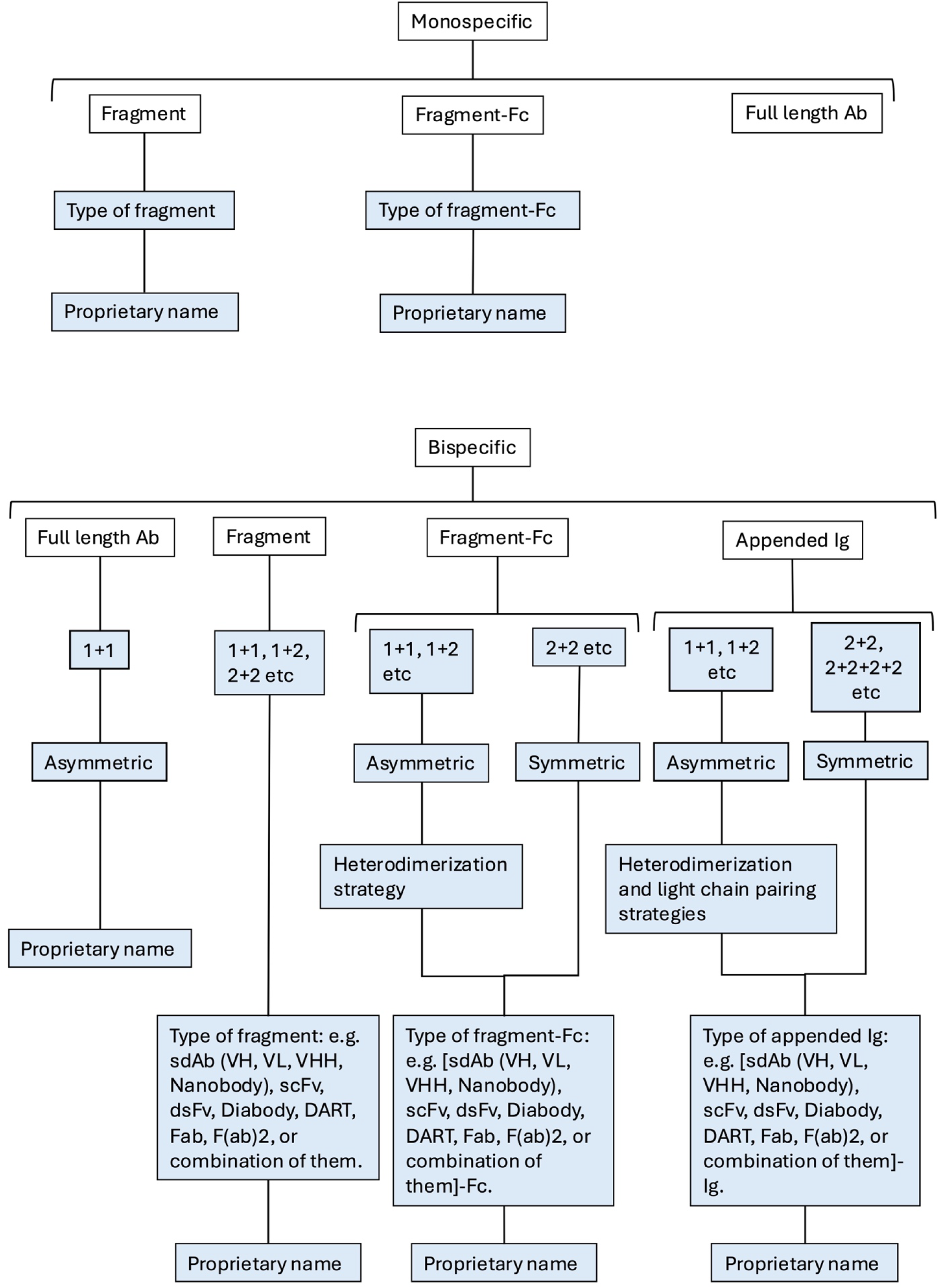
Schematic of detailed format classification system. The schematic represents the rationale behind the additional information we provide on the format. When available, we provide information on the type of format (e.g. type of fragment, type of fragment-Fc, type of fragment-Ig) and proprietary name. For bispecific antibodies, when available and if applicable, we also provide information on valency, symmetry, and heterodimerization and light chain pairing strategies.

**Supp. Figure S3.**
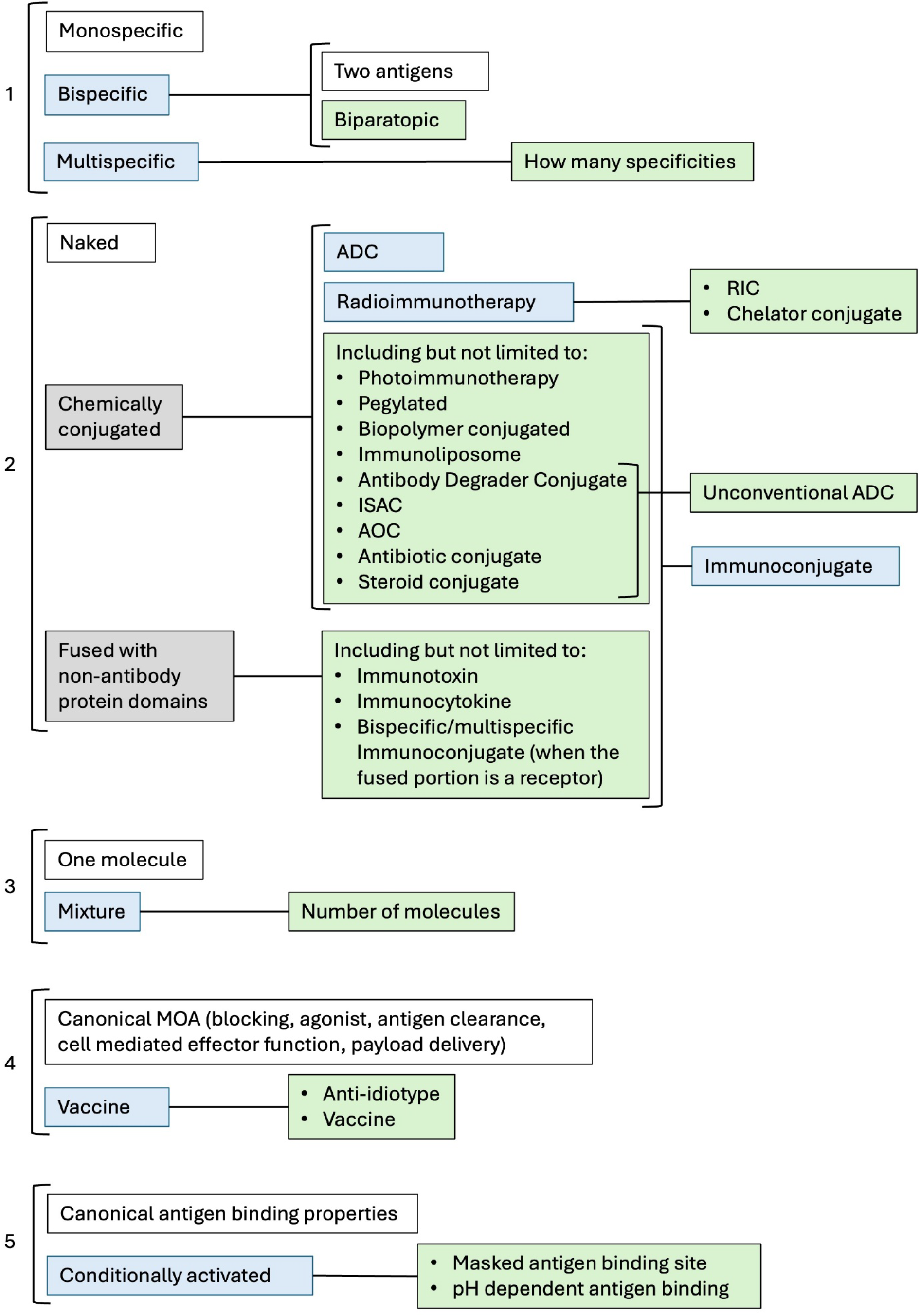
Schematic of general molecular classification system. To describe antibody general molecular category, we considered five possible features that can describe the molecule: 1) specificity, 2) conjugation status, 3) one molecule or mixture drug composition, 4) mechanism of action (MOA), 5) antigen binding properties. Blue boxes represent the broad variables displayed in the “General molecular category” category. Green boxes represent additional variables that describe the molecules in detail. Grey boxes are represented in the schematic to better describe immunoconjugates, but are not displayed in the database. Unless stated otherwise with the variables described in the blue and green boxes, we consider the molecules to be naked, with canonical MOA (blocking, agonist, antigen clearance, cell mediated effector function, payload delivery) and canonical antigen binding properties. Naked monospecific antibodies are labelled as “Naked monospecific”, and, unless stated otherwise, we consider them to have canonical MOA and canonical antigen binding properties. Each molecule can be described by one or more of the variables described above.

**Supp. Figure S4.**
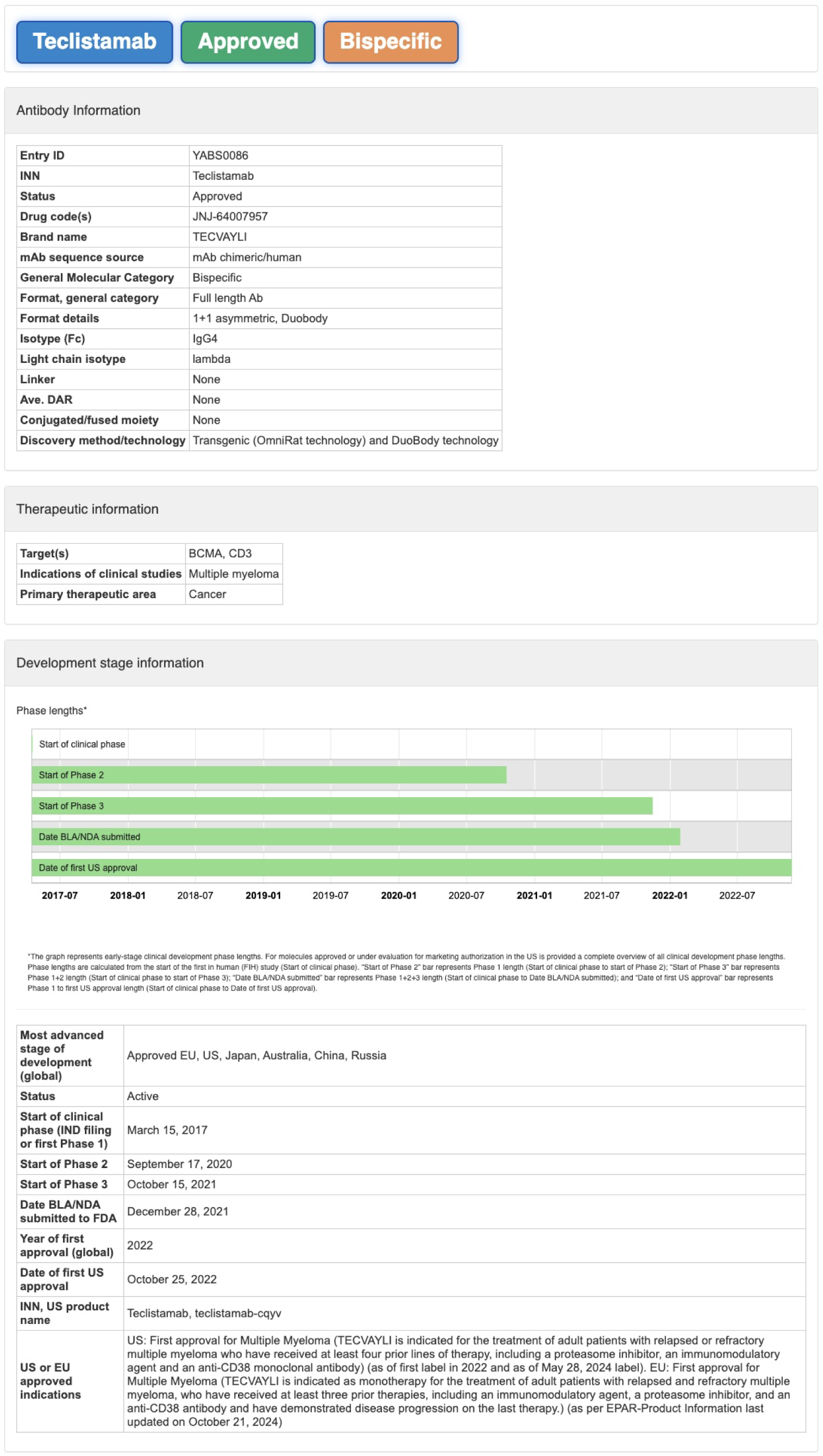
Example of a molecule dedicated page, top part. Each molecule dedicated page has three labels at the top showing INN (when available) or drug code, general status and general molecular category. Information is then grouped in sections, as in the example provided, showing antibody information (including names, drug codes, and molecular characteristics); therapeutic information (including target, indications and primary therapeutic area); and development stage information (including most advanced stage of development, status, key milestone dates of clinical development and, where applicable, US and/or EU approved indications). The development stage information section includes a graphical overview of early-stage clinical development phase lengths, and for antibodies in regulatory review/approved in US, a complete overview of all clinical development phase lengths. The other information provided in the molecule dedicated page is shown in **Supplemental Figure S5**.

**Supp. Figure S5.**
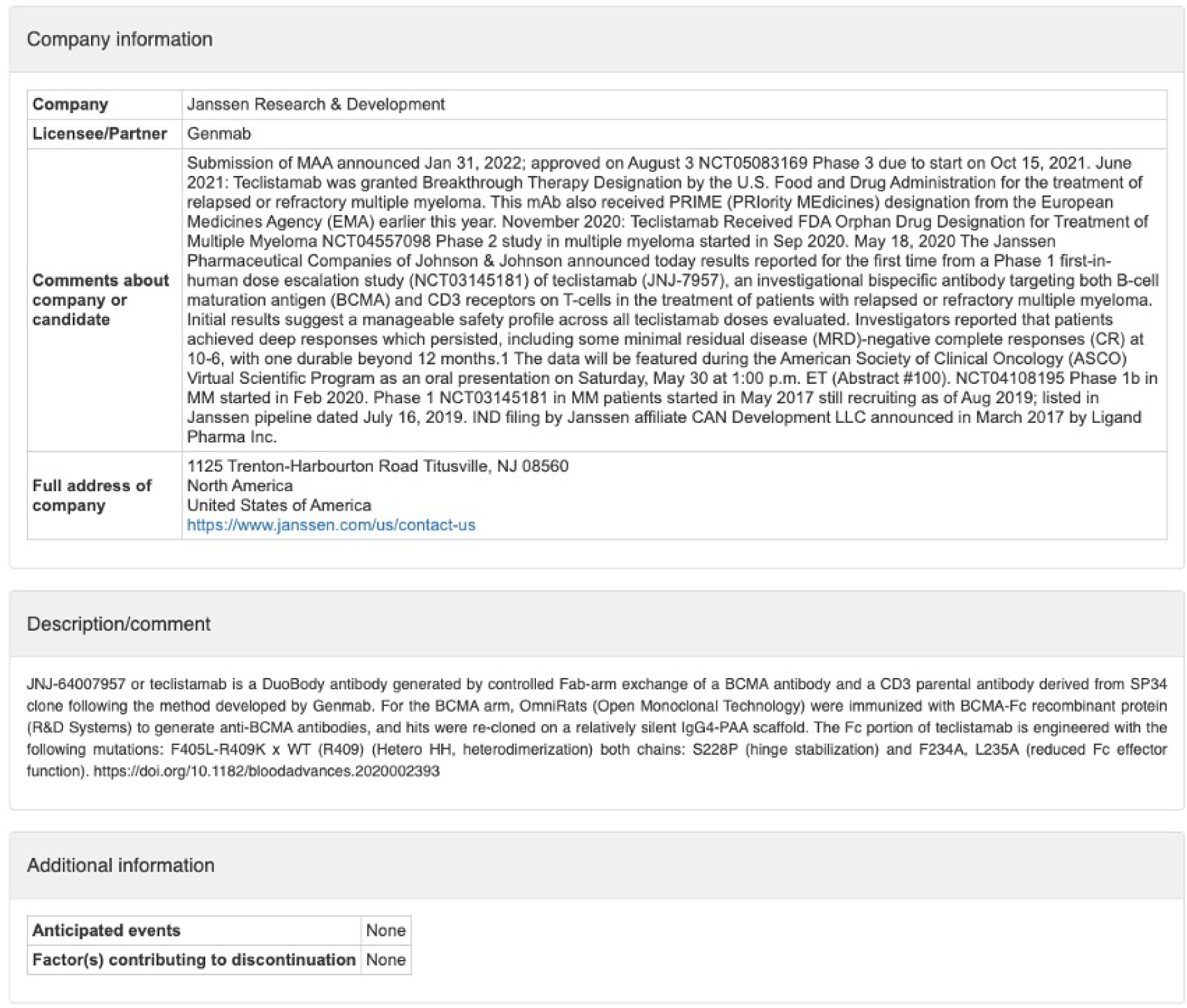
Example of a molecule dedicated page, bottom part. Each molecule dedicated page has the features described in **Supplemental Figure S4** followed by the sections shown in the example provided: company information (including names of the company and eventual licensee/partner, comments about company or candidate molecule and full address of the company); description/comment (including, when available, detailed information on the molecule and supporting references); additional information (including anticipated events, when available, and factors contributing to discontinuation, when applicable and if available).

**Supp. Figure S6.**
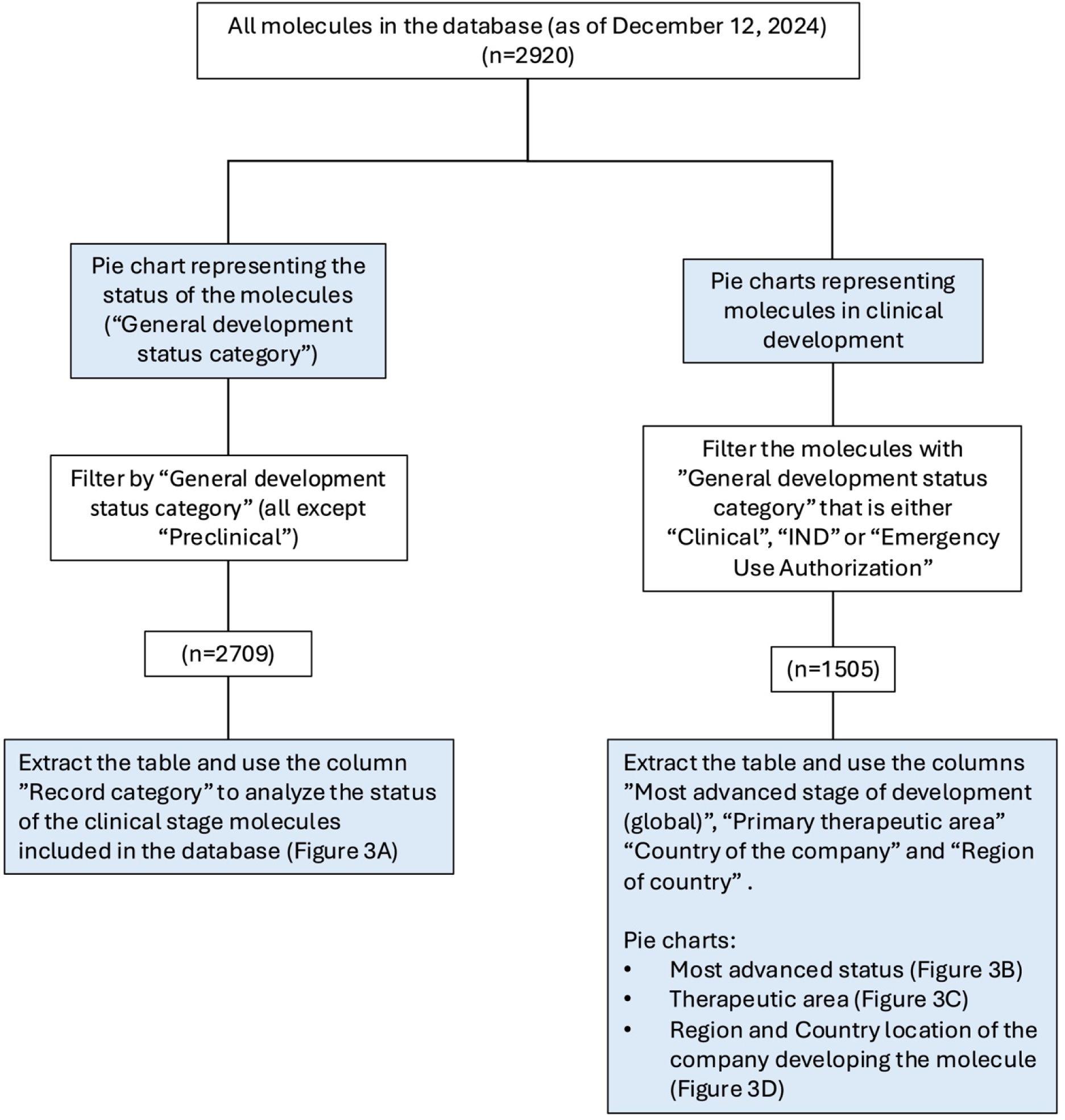
Analysis pipeline for Figure 3. Filtering criteria and selected columns of the exported table used for analyzing the data included in the graphs for Figure 3.

**Supp. Figure S7.**
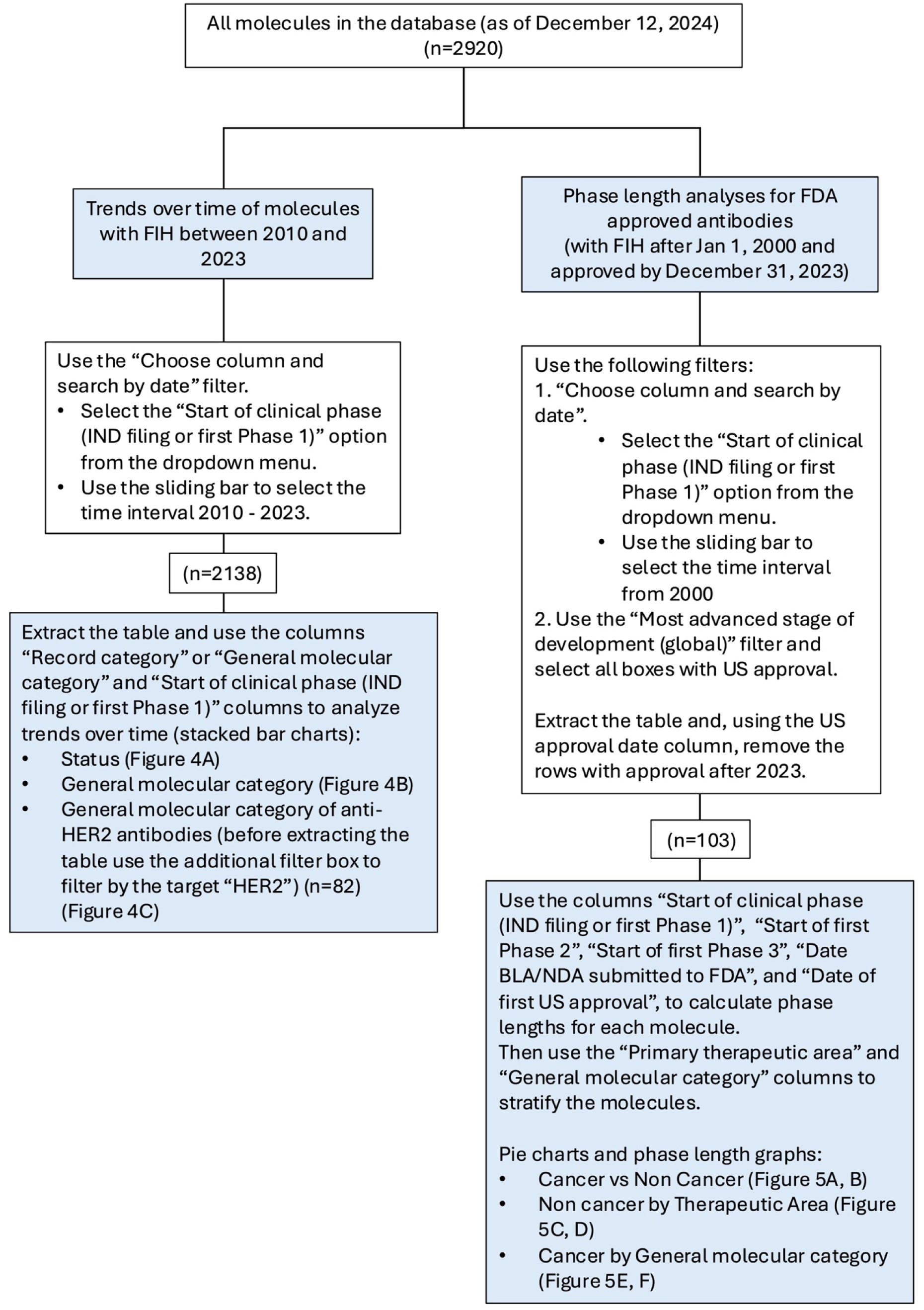
Analysis pipeline for Figures 4 and 5. Filtering criteria and selected columns of the exported table used for analyzing the data included in the graphs for Figure 4 **and 5**.

**Supp. Table S1.**
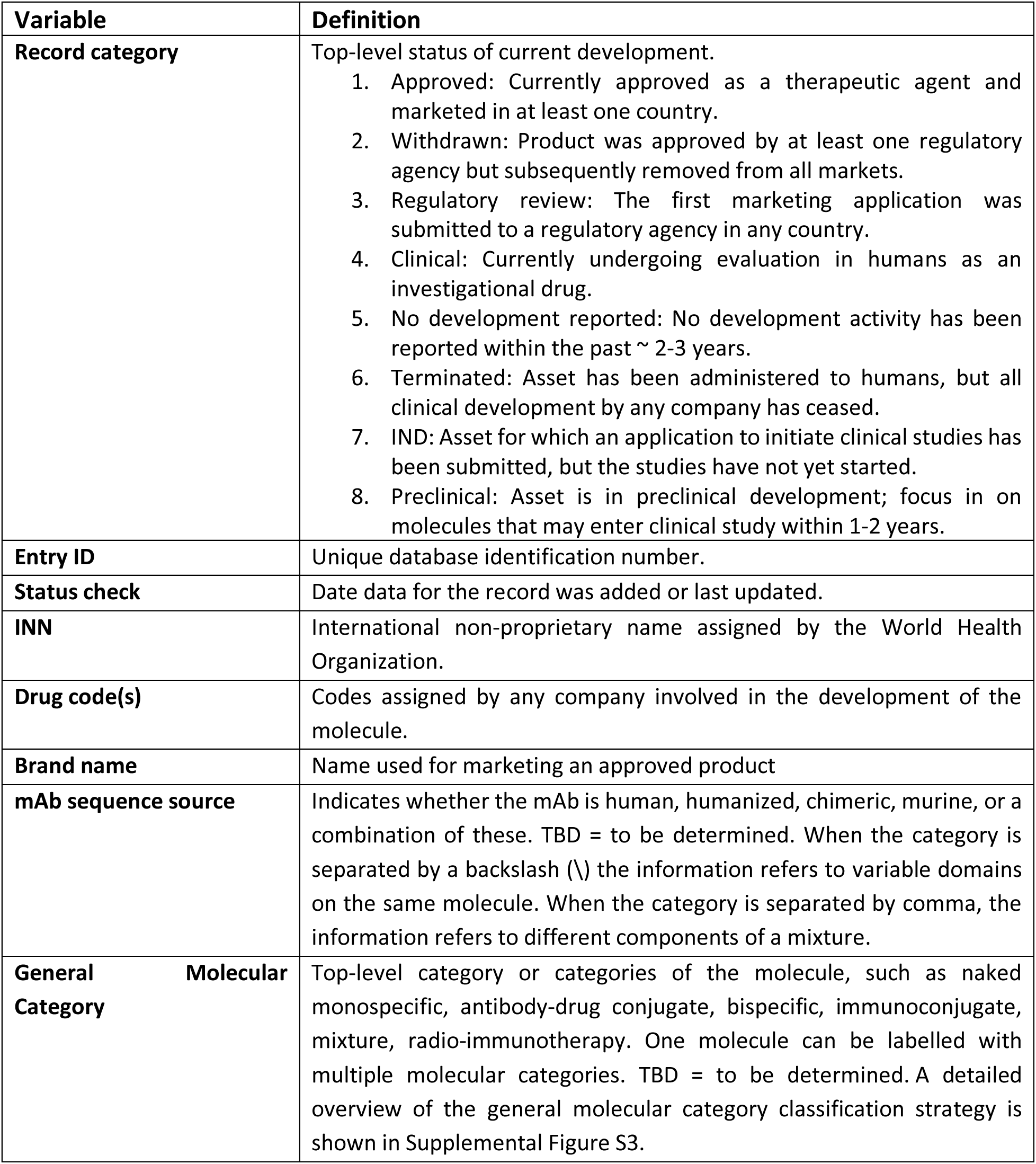
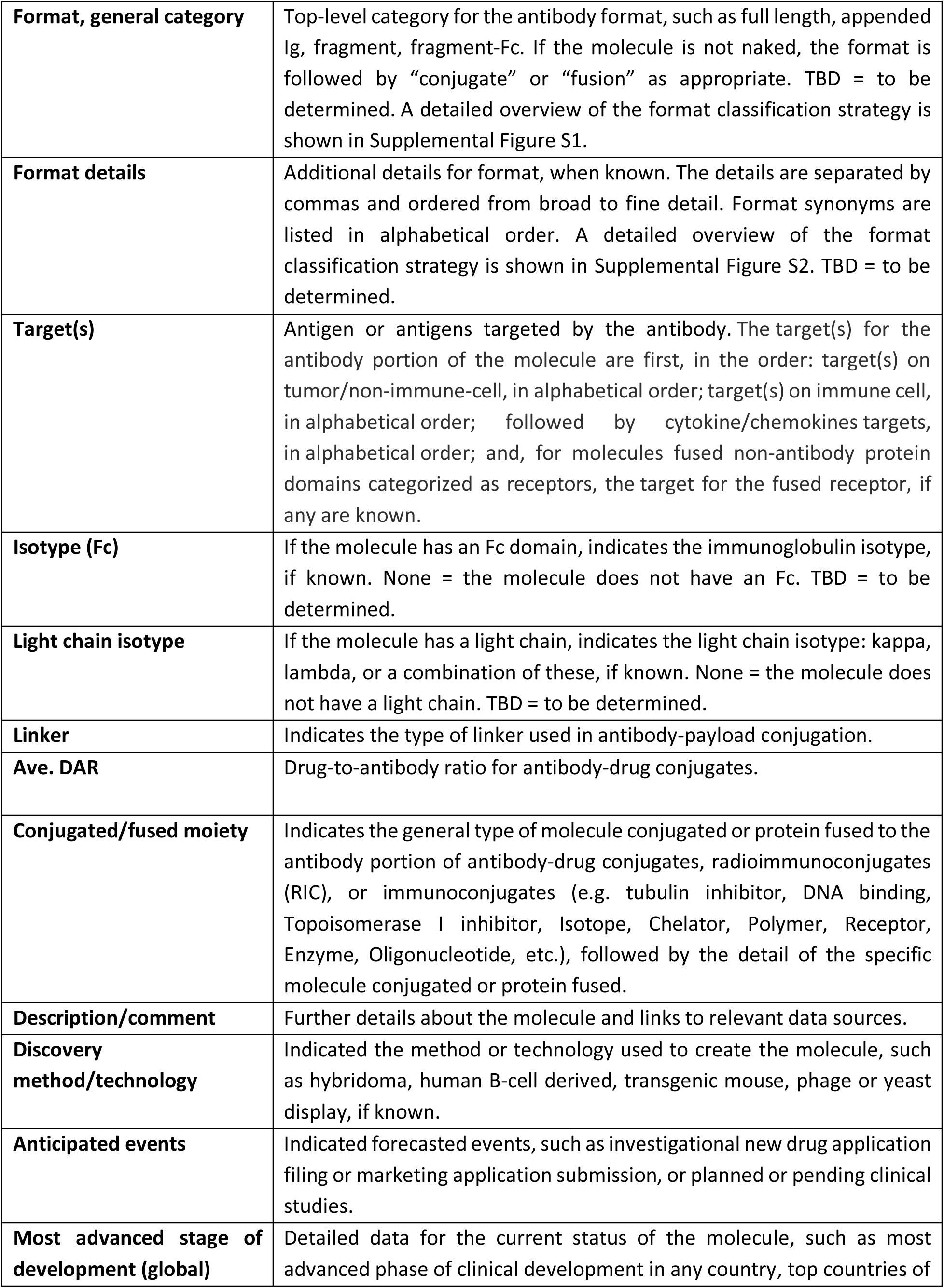
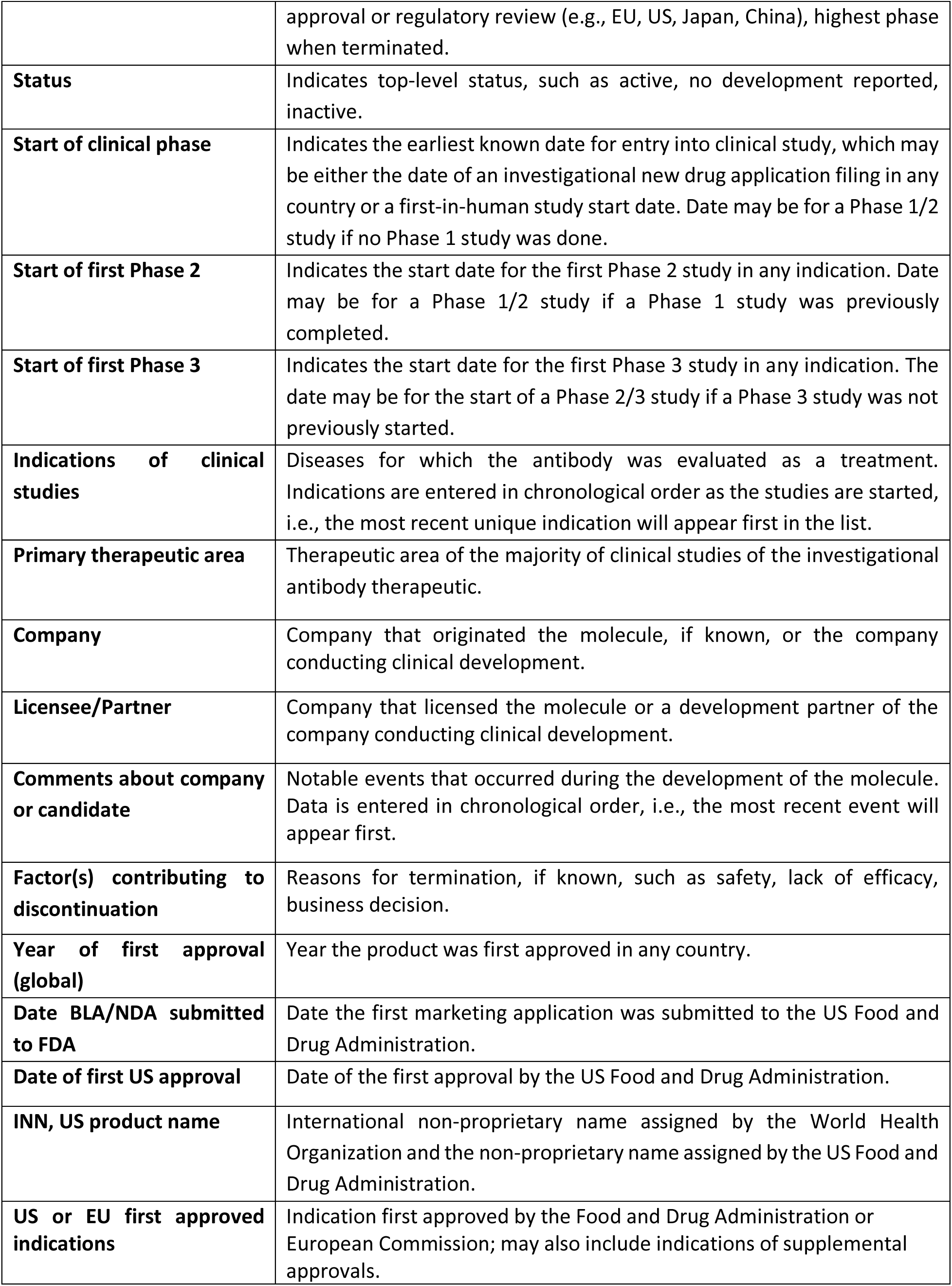
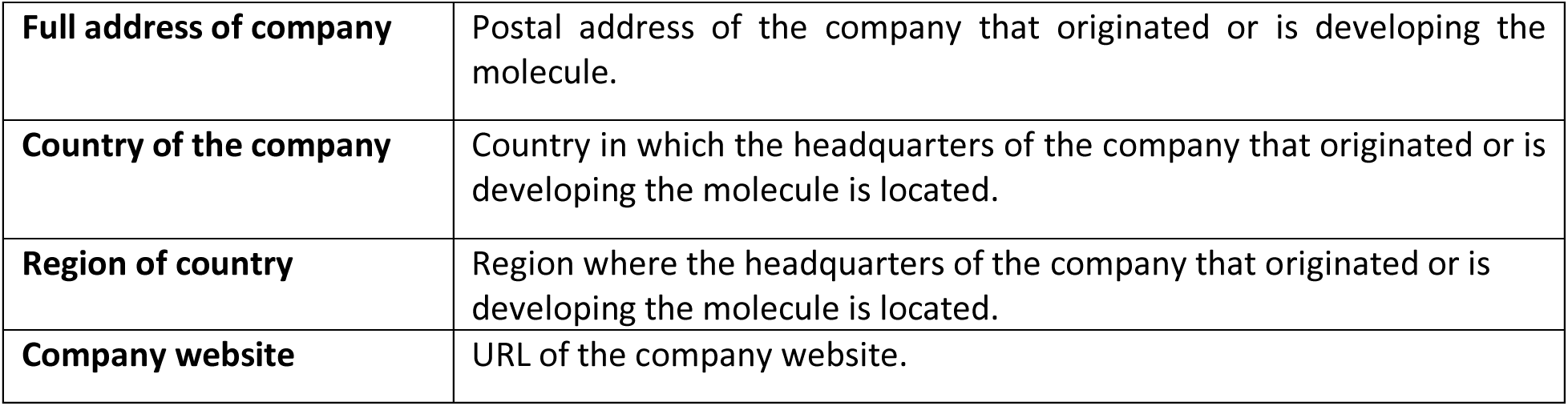
Database variables and definitions.

## References

1. Crescioli S, Jatiani S, Moise L. With great power, comes great responsibility: the importance of broadly measuring Fc-mediated effector function early in the antibody development process. MAbs. 2025;17(1). doi: 10.1080/19420862.2025.2453515.

2. Wilkinson I, Hale G. Systematic analysis of the varied designs of 819 therapeutic antibodies and Fc fusion proteins assigned international nonproprietary names. MAbs. 2022;14(1). doi: 10.1080/19420862.2022.2123299.

3. Sanou G, Manso T, Todorov K, Giudicelli V, Duroux P, Kossida S. IMGT/mAb-KG: the knowledge graph for therapeutic monoclonal antibodies. Front Immunol 2024;15. doi: 10.3389/fimmu.2024.1393839.

4. Kaplon H, Crescioli S, Chenoweth A, Visweswaraiah J, Reichert JM. Antibodies to watch in 2023. MAbs. 2023;15(1). doi.org/10.1080/19420862.2022.2153410.

5. Crescioli S, Kaplon H, Chenoweth A, Wang L, Visweswaraiah J, Reichert JM. Antibodies to watch in 2024. MAbs. 2024;16(1). doi.org/10.1080/19420862.2023.2297450.

6. Crescioli S, Kaplon H, Wang L, Visweswaraiah J, Kapoor V, Reichert JM. Antibodies to watch in 2025. Mabs 2025; 17(1). doi.org/10.1080/19420862.2024.2443538.

7. Amash A, Volkers G, Farber P, Griffin D, Davison KS, Goodman A, Tonikian R, Yamniuk A, Barnhart B, Jacobs T. Developability considerations for bispecific and multispecific antibodies. MAbs. 2024;16(1). doi: 10.1080/19420862.2024.2394229.

8. Klein C, Brinkmann U, Reichert JM, Kontermann RE. The present and future of bispecific antibodies for cancer therapy. Nat Rev Drug Discov. 2024;23(4):301–319.

9. Labrijn AF, Janmaat ML, Reichert JM, Parren PWHI. Bispecific antibodies: a mechanistic review of the pipeline. Nat Rev Drug Discov. 2019;18(8):585–608. doi: 10.1038/s41573-019-0028-1.

10. Brinkmann U, Kontermann RE. The making of bispecific antibodies. MAbs. 2017;9(2):182–212. doi: 10.1080/19420862.2016.1268307.

11. Dumontet C, Reichert JM, Senter PD, Lambert JM, Beck A. Antibody-drug conjugates come of age in oncology. Nat Rev Drug Discov. 2023;22(8):641–661.

12. Kaplon H, Reichert JM. Antibodies to watch in 2019. mAbs 2018, 11(2), 219–238. doi.org/10.1080/19420862.2018.1556465.

13. Knox C, Wilson M, Klinger C M, Franklin M, Oler E, Wilson A, Pon A, Cox J, Chin NEL, Strawbridge S A, et al. DrugBank 6.0: the DrugBank Knowledgebase for 2024. Nucleic Acids Res. 2024, 52(D1), D1265–D1275.

14. Raybould MIJ, Marks C, Lewis AP, Shi J, Bujotzek A, Taddese B, Deane CM. Thera-SAbDab: the Therapeutic Structural Antibody Database. Nucleic Acids Res. 2020;48(D1):D383–D388. doi: 10.1093/nar/gkz827.

15. Rawat P, Sharma D, Prabakaran R, Ridha F, Mohkhedkar M, Janakiraman V, Gromiha MM. Ab-CoV: a curated database for binding affinity and neutralization profiles of coronavirus-related antibodies. Bioinformatics 2022; 38:4051–2.

16. Rawat P, Prabakaran R, Sakthivel R, Mary Thangakani A, Kumar S, Gromiha MM. CPAD 2.0: a repository of curated experimental data on aggregating proteins and peptides. Amyloid 2020; 27:128–33.

17. Khan A, Cowen-Rivers AI, Grosnit A, Deik DG, Robert PA, Greiff V, Smorodina E, Rawat P, Akbar R, Dreczkowski K, et al. Toward real-world automated antibody design with combinatorial Bayesian optimization. Cell Rep Methods. 2023;3(1):100374. doi: 10.1016/j.crmeth.2022.100374.

18. Akbar R, Bashour H, Rawat P, Robert PA, Smorodina E, Cotet TS, Flem-Karlsen K, Frank R, Mehta BB, Vu MH, et al. Progress and challenges for the machine learning-based design of fit-for-purpose monoclonal antibodies. MAbs. 2022;14(1). doi: 10.1080/19420862.2021.2008790.

